# Differences in developmental potential predict the contrasting patterns of dental diversification in characiform and cypriniform fishes

**DOI:** 10.1101/2020.07.26.221986

**Authors:** David Jandzik, David W Stock

## Abstract

Morphological diversification during adaptive radiation may depend on factors external or internal to the lineage. We provide evidence for the latter in characiform fishes (tetras and piranhas), which exhibit extensive dental diversity. Phylogenetic character mapping supported regain of lost teeth as contributing to this diversity. To test for latent potential for dentition that would facilitate its evolutionary expansion, we overexpressed a tooth initiation signal, the tumor necrosis factor pathway ligand ectodysplasin, in a model characiform, the Mexican Tetra (*Astyanax mexicanus*). This manipulation resulted in extensive ectopic dentition, in contrast to its previously-reported limited effect in the Zebrafish (*Danio rerio*). Tooth location in the Order Cypriniformes, to which the Zebrafish belongs, is much more restricted than in characiforms, a pattern that may be explained by differences in the retention of ancestral developmental potential. Our results suggest that differences in evolvability between lineages may lead to contrasting patterns of diversification.

## Introduction

The morphological diversity present in a clade of organisms is influenced both by the environments encountered by the included species, as well as their evolvability – the capacity to generate adaptive variation (Wagner & Altenberg 1996, Gerhart & Kirshner 2003; Hendrikse *et al.* 2007, Pigliucci 2008, Erwin 2017). One manner in which evolvability might be manifest is the biasing of phenotypic variants toward those that were adaptive in the past (Watson *et al.* 2014, Watson & Szathmáry 2016). An example is provided by the retention and re-expression of “ancestral developmental potential” for a specific caste morphology in the evolution of ants (Rajakumar *et al.* 2012). The degree to which such potential differs among clades and whether these differences are responsible for differing patterns of morphological diversification remains largely unknown, however.

In a previous study (Aigler *et al.* 2014), we used overexpression of a tooth initiation signal encoded by the *ectodysplasin* (*eda*) gene to show that the Zebrafish (*Danio rerio*), a species with highly reduced dentition, retains limited potential to re-express teeth in ancestral locations. This limited potential is consistent with the pattern of dental diversification of the order Cypriniformes, to which the Zebrafish belongs. Cypriniform fishes, which include carps, loaches, minnows, suckers and over 4200 species, are dominant elements of the freshwater fish faunas of North America, Africa and Eurasia (Nelson *et al.* 2016). Despite exploiting a diversity of food sources, ranging from detritus to plants to insects to other fishes (Howes 1991), teeth in this group are restricted to a single pair of bones (fifth ceratobranchials) in the lower posterior pharynx (Stock 2007).

In contrast to the limited extent and evolutionary conservatism of tooth location in the Cypriniformes, the members of the related order Characiformes (tetras, piranhas, and relatives) generally exhibit a more extensive dentition; in addition, considerable variation in tooth location exists among species. The order Characiformes is actually smaller than the Cypriniformes (approximately 2300 species), exhibits a comparable diversity of diets (Guisande *et al.* 2012) and while co-occurring with cypriniforms in North America and Africa, is a dominant element of the freshwater fish fauna of South America, which lacks cypriniforms (Nelson *et al.* 2016). Teeth in characiforms may be found on marginal bones of the oral jaws (including their surfaces outside of the mouth), bones of the palate, paired bones and gill rakers of the upper and lower pharynx, and midline bones of the floor of the mouth and pharynx (Fink & Fink 1981; Novakowski *et al.* 2004; Oyakawa & Mattox 2009, Roberts 1969; 1973; Toledo-Piza 2000; 2007; Weitzman 1962; Weitzman & Fink 1985).

In the present study, we tested the hypothesis that the greater variability of tooth location in characiforms relative to cypriniforms is the result of a difference in evolvability, and specifically a greater retention of ancestral potential for dentition in the former group. This hypothesis is based on the commonly-held view that the ancestral condition of dentition in bony fishes consisted of teeth on virtually all of the bones lining the oral and pharyngeal cavities, as can be seen in the extant bowfin (*Amia calva*) (Grande & Bemis 1998; Stock 2001). The toothless bones of the mouth and pharynx of any characiform (or cypriniform) species therefore bore teeth at some point in its ancestry. An alternative to our hypothesis on the cause of variability in tooth location in characiforms that does not involve retention of ancestral potential is that variability in tooth location among species arises simply from loss of teeth from the extensive dentition of the common ancestor of this group. We used phylogenetic character mapping to show that while teeth have indeed been lost within the characiforms in this manner, there have also been instances of re-expansion of the dentition, a phenomenon that might involve the realization of latent developmental potential. We next tested for the existence of such potential in the characiform Mexican Blind Cave Tetra (*Astyanax mexicanus*) by overexpression of ectodysplasin. We found that such expression was capable of greatly expanding both the larval and adult dentitions of this species. Bones bearing ectopic teeth included several that have regained lost teeth in characiform evolution, as well as others from which teeth are absent in all characiforms but have been regained in other lineages outside of this group. In addition to supporting our specific hypothesis that dental evolution in characiforms has resulted from the realization of retained latent potential for dentition, our results suggest that differences in morphological outcomes in related groups radiating in similar environments may result from differences in evolvability.

## Results

### Distribution of teeth in the Characiformes

Because of the absence of a concise summary of all of the bones that may bear teeth in characiforms, we surveyed the osteological and taxonomic literature of this group (Weitzman 1962; Roberts 1969; 1973; Fink & Fink 1981; Weitzman & Fink 1985; Toledo-Piza 2000; 2007; Novakowski *et al.* 2004; Oyakawa & Mattox 2009) to produce our own (Fig. 1J, L). Teeth may be found on all of the bones of the jaw margins – the premaxillaries and maxillaries of the upper jaw and the dentaries of the lower jaw (Fig. 1B-D). In some genera, such as *Tyttocharax* and *Roeboides*, teeth are present on the surfaces of these bones that extend outside of the mouth (Fig. 1G).

**Figure 1.**
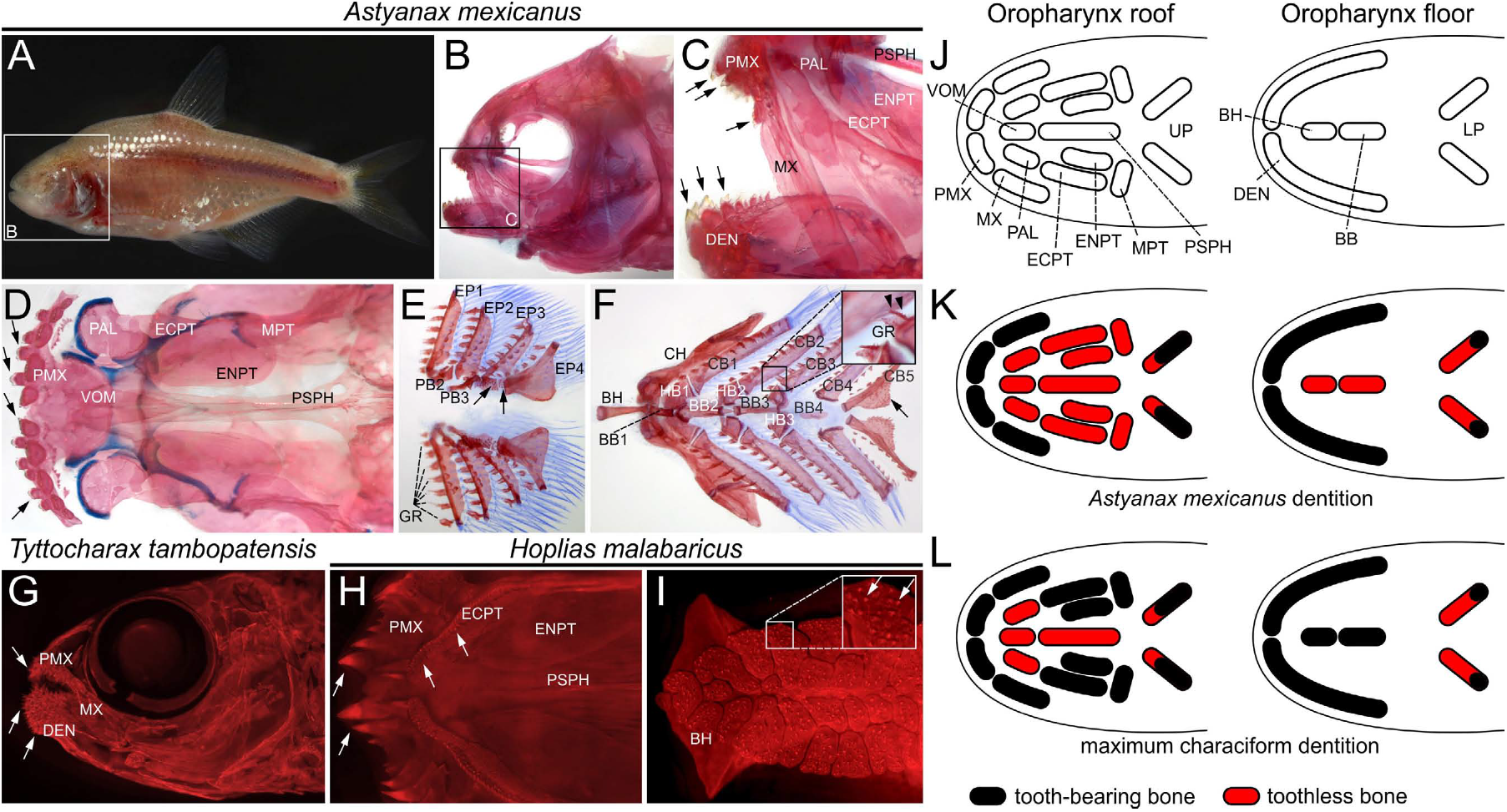
Distribution of teeth in wild type *Astyanax mexicanus* and other members of the Order Characiformes. A) Lateral view of adult cave-dwelling morph of *A*. *mexicanus*. B-F) Alizarin red- and Alcian blue-stained intact (B, C) and dissected (D-F) head skeletons of adult *A*. *mexicanus*. Lateral view of jaw margins (B, C), ventral view of palate (D), ventral view of dorsal gill arches (E) and dorsal view of ventral hyoid and gill arches (E, F). Teeth (arrows) are present on premaxillary (C, D), maxillary (C) and dentary (C) bones (C), of the jaw margins, are absent from the palate (D), and are present on tooth plates supported by pharyngobranchials and epibranchials of the dorsal gill arches (E), as well as the fifth ceratobrancials of the lower gill arches (F). Teeth are additionally present on gill rakers attached to epibranchials (E), hypobranchials and ceratobranchials (arrowheads in F). G) Lateral view of cleared and alizarin-stained head skeleton of *Tyttocharax tambopatensis* (Characiformes: Characidae) showing teeth (arrows) outside of the mouth on the premaxillary and dentary bones. H-I) Cleared and alizarin-stained palate (H) and basihyal (I) of *Hoplias malabaricus* (Characiformes: Erythrinidae). Anterior arrows in H indicate premaxillary teeth and middle and posterior arrows indicate accessory ectopterygoid and ectopterygoid teeth, respectively. Arrows in I indicate fine teeth attached to tooth plates supported by the basihyal (“tongue”). J-L) Schematic of bones lining the roof (left drawing) and floor (right drawing) of the oropharynx modelled after Figure 16 of Gosline (1971). Gill arches have been simplified as a single element with anterior and posterior ends corresponding to the position of individual arches along the anterior-posterior axis. Bones that bear teeth in *A*. *mexicanus* (K) and may bear teeth when the entire Order Characiformes is considered (L) are indicated in black, and bones without teeth in red. Abbreviations: BB, basibranchial; BH, basihyal; CB, ceratobranchial; CH, ceratohyal; DEN, dentary; ECPT, ectopterygoid; ENPT, endopterygoid; EP, epibranchial; GR, gill raker; HB, hypobranchial; MPT, metapterygoid; MX, maxillary; PAL, palatine; PB, pharyngobranchial; PMX, premaxillary; PSPH, parasphenoid; UP, upper pharyngeal elements; LP, lower pharyngeal elements; VOM, vomer.

The roof of the mouth (loosely the palate) of teleost fishes is lined medially by bones of the ventral braincase and laterally by bones comprising the hyopalatine arch or suspensorium (Hilton 2011). In characiforms, palatal teeth are limited to the suspensorium, and may be present on the ectopterygoids, endopterygoids and metapterygoids (Figure 1H, J-L). An additional tooth plate anterior to the ectopterygoid that is present in members of the families Erythrinidae (Fig. 1H) and Hepsetidae has been considered neomorphic (an accessory ectopterygoid) rather than a dermopalatine, which is absent in characiforms but occupies a similar position in some teleosts (Fig 1H) (Roberts 1969, 1973, Fink & Fink 1981).

As in cypriniforms, teeth may be found in the lower pharynx on the fifth ceratobranchials (last gill arch) (Fig 1F, J-L). Unlike cypriniforms, characiforms may have teeth on upper pharyngeal tooth plates supported by the second and third pharyngobranchials and third and fourth epibranchials (Fig. 1E, J-L). Teeth may also be found on gill rakers attached to all five gill arches (Fig. 1F). In the midline of the mouth and pharynx, teeth may be found (rarely) on tooth plates attached to the basihyal (“tongue”) and basibranchials (Fig 1I-J, L).

### Loss and reappearance of teeth in the evolution of characiforms

We next searched for evidence that teeth had reappeared during the evolution of the Characiformes. Teeth on the metapterygoid bones of the suspensorium are extremely rare in ray-finned fishes, being found only in the non-teleost families Amiidae (the Bowfin) and Polypteridae (bichirs), as well as the characiform genera *Hydrolycus* and *Raphiodon* of the Cynodontinae (Toledo-Piza 2000). This subfamily is nested within the Characiformes in molecular (Oliveira *et al.* 2011; Arcila *et al.* 2017; Betancur-R *et al.* 2019), morphological (Mirande 2009, 2010) and combined (Mirande 2019) phylogenies, providing strong support for the reappearance of these teeth after an absence of 200-300 million years (Irisarri *et al.* 2017; Hughes *et al.* 2018). Mirande (2009, 2010) compiled a morphological dataset for 160 characiform species that allows mapping the presence or absence of teeth on the premaxillaries outside of the mouth, maxillaries, ectopterygoids, endopterygoids, fourth basibranchial, gill rakers, fifth ceratobranchials, and pharyngobranchials (third, fourth, and fifth) on his phylogeny (Fig. S1, S2). Premaxillary teeth outside of the mouth are not present in non-teleostean ray-finned fishes (Nelson *et al.* 2016) but appeared in multiple characiform lineages. Maxillary teeth are reconstructed as having appeared within the Characiformes, but use of alternative outgroups would likely change this interpretation (Stock 2007). Reappearance of teeth within the Characiformes is supported for ectopterygoid, endopterygoid, and basibranchial bones, as well as gill rakers. To test the robustness of a subset of these results, we mapped presence and absence of ectopterygoid and endopterygoid teeth onto the molecular phylogeny of Oliveira *et al.* (2011) (Fig. 2). We chose these teeth because of the necessity of compiling a character matrix (Table S1; Supplementary References) for taxa not present in Mirande’s (2009) analysis, which was facilitated by the fact that these teeth are commonly mentioned in taxonomic studies of characiform species. Our analysis suggested that ectopterygoid teeth were present in the common ancestor of characiforms and were regained after loss four times within the group (Fig. 2A, S3A). Endopterygoid teeth were reconstructed as absent in the characiform common ancestor and were gained five times within the group (Fig. 2B, S3B). Reappearance of ectopterygoid and endopterygoid teeth during characiform evolution was also supported by a similar analysis using the phylogeny of Betancur-R *et al.* (2019) (Fig. S3C, D; Table S2; Supplementary References).

**Figure 2.**
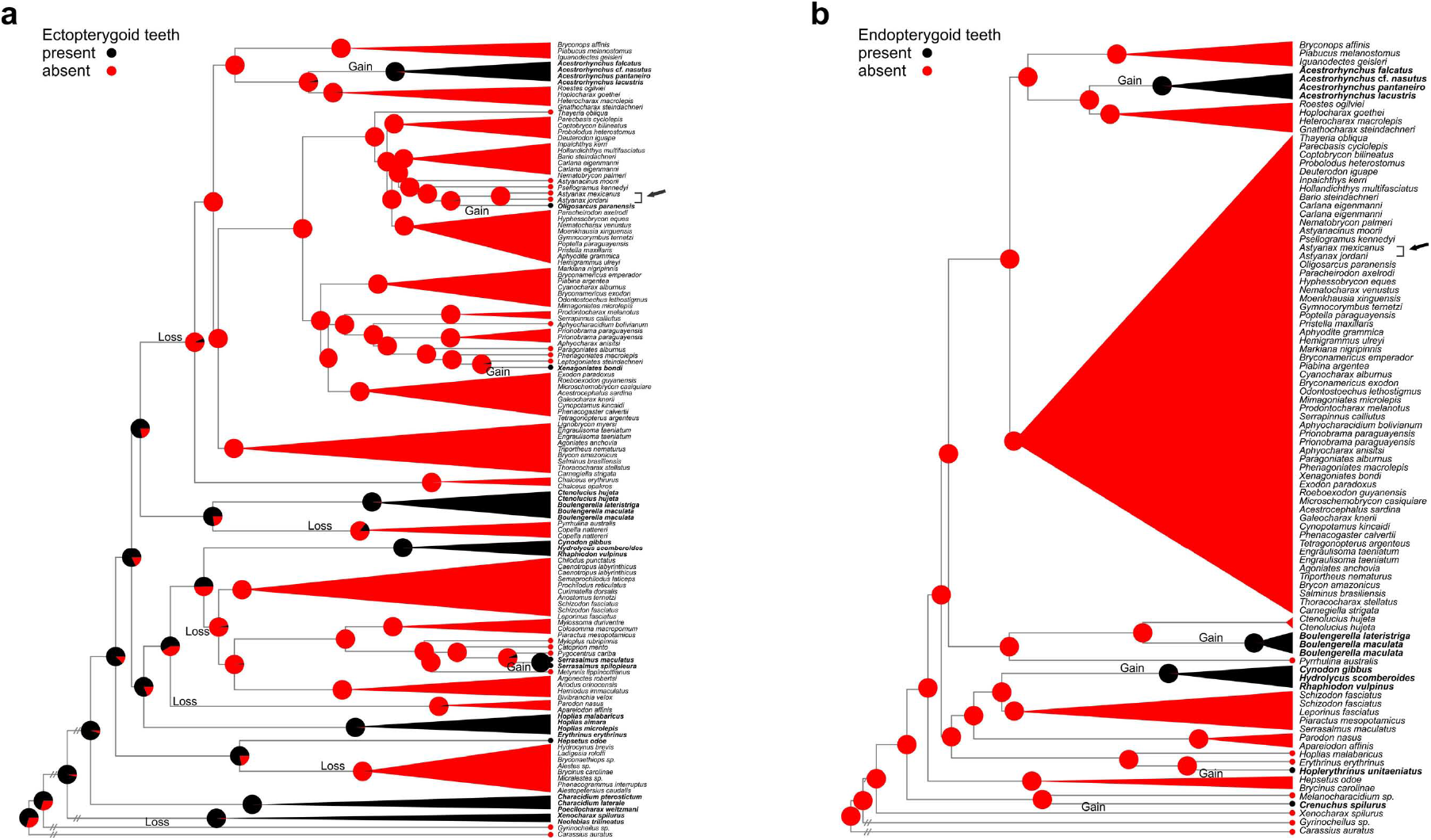
Teeth on the ectopterygoid and endopterygoid bones have been re-gained following loss in multiple characiform lineages. Maximum likelihood ancestral state reconstruction of presence (black) or absence (red) of teeth on the ectopterygoid (A) and endopterygoid (B) bones. Tree topologies and branch lengths are from Oliveira *et al.* (2011) and character states were compiled from the literature (Supplementary References and Table S1). Hashmarks indicate branches not drawn to scale. Pie charts at nodes represent the relative probabilities of each character state. Clades with no internal character change were collapsed; full versions of the tree are presented in Figure S3(A, B) and reconstructions based on alternative phylogenies in Figures S1 and S3(C, D). *Carassius auratus* and *Gyrinocheilus* sp. are outgroups within the Order Cypriniformes, while the position of *A*. *mexicanus* (and its cave morph, sometimes designated *A*. *jordani*) is indicated with an arrow. Ectopterygoid teeth are reconstructed with highest probability as being present in the characiform common ancestor, with six losses and four re-gains occurring within the group. Endopterygoid teeth (lacking in the characiform common ancestor) were gained at least five times in the Characiformes.

### Larval dentition of eda-overexpressing A. mexicanus

The developmental genetic basis of the loss and reappearance of teeth in characiform evolution remains unknown. A candidate cause is modification of the ectodysplasin signaling pathway, which has been shown in the Zebrafish to be both necessary for tooth development (Harris *et al.* 2008) as well as sufficient for expanding tooth-bearing locations (Aigler *et al.* 2014). We tested the ability of altered *eda* signaling to induce ectopic teeth in a model characiform species, the Mexican Blind Cave Tetra, *Astyanax mexicanus* (Jeffery 2009; Casane & Rétaux 2016), by injection of an *eda* overexpression construct into one-celled embryos. Injection of a similar construct in the Zebrafish expanded dentition along the dorsal-ventral axis, but not along the anterior-posterior axis (Aigler *et al.* 2014). Specifically, teeth in wild type zebrafish are found only in the posterior ventral pharynx, while overexpression of *eda*-induced ectopic teeth in this location, as well as the posterior dorsal pharynx.

The wildtype dentition of *A*. *mexicanus* is similar to that of numerous characiforms, with teeth being found on the premaxillary, maxillary and dentary bones of the oral jaw margins, the fifth ceratobranchial bones of the lower pharynx, and dorsal pharyngeal tooth plates attached to the second and third pharyngobranchials, as well as the third and fourth epibranchials (Figure 1B-F, J-K) (Valdéz-Moreno & Contreras-Balderas 2003). In addition, teeth are present on gill rakers attached to dorsal and ventral elements of the anterior four gill arches, as well as the ventral fifth ceratobranchials (Atukorala & Franz-Odendaal 2014). In our initial injections of the *eda*-overexpression construct, we examined larvae stained for calcified structures with alizarin red at 6 days post-fertilization (dpf). In wild type larvae of this age, teeth are limited to the premaxillary and dentary bones of the oral jaws, the fifth ceratobranchials and the posterior-most upper pharyngeal toothplate (Trapani *et al.* 2005; Atukorala & Franz-Odendaal 2014), *i*.*e*. dorsally and ventrally at the anterior and posterior margins of the oropharyngeal cavity (Fig. 3A-H). We found ectopic teeth in 47 of 397 (11.8%) larvae surviving to 6 dpf following injection with the *eda*-overexpression construct and none of the 33 surviving control larvae injected with a similar construct for expressing green fluorescent protein (*gfp*) (p = 0.0372, Fisher’s exact test). In contrast to our previous results with the zebrafish (Aigler *et al.* 2014), we found that *eda* overexpression was capable of expanding the dentition into the central part of the oropharyngeal cavity, including laterally on anterior ceratobranchials and medially in the ventral basibranchial area (Fig. 3E-H).

**Figure 3.**
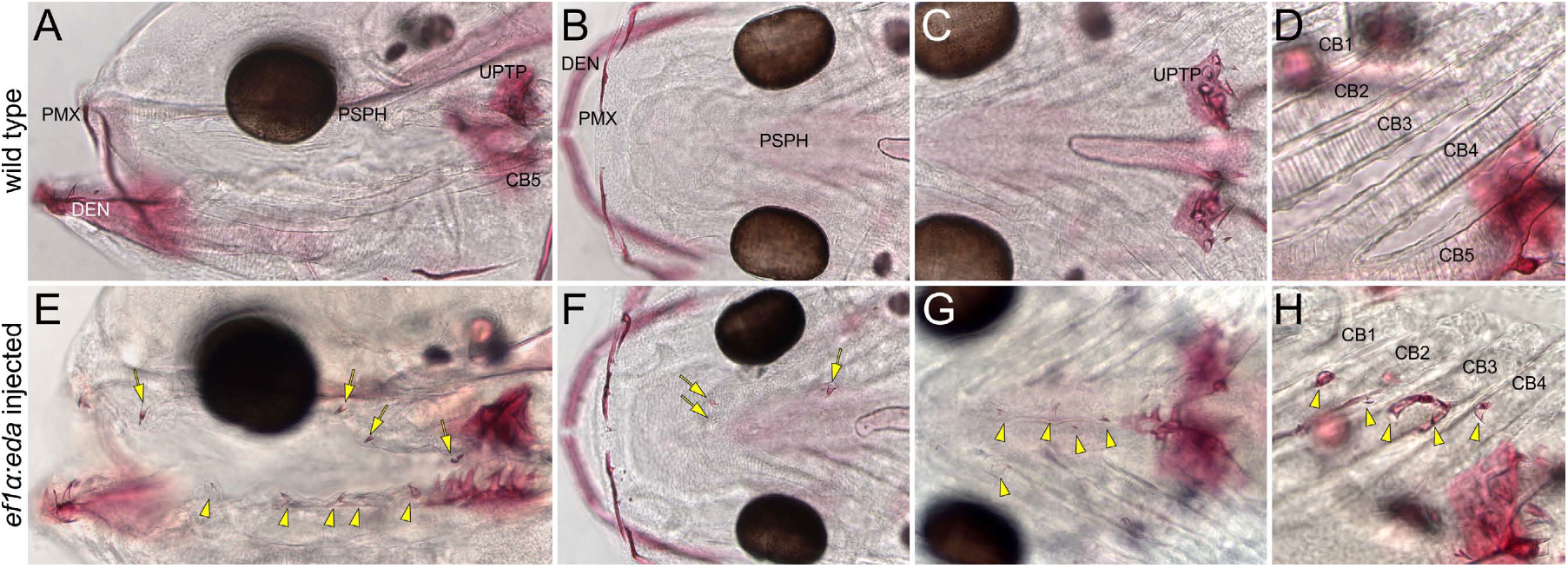
Ectopic expression of eda expands the larval dentition of *A*. *mexicanus* into the central oropharynx. Cleared and alizarin-stained wild type (A-D) and *pEF1α:EDA*-injected larvae (E-H) at six dpf in lateral (A, E), dorsal (B-D, F-G) and ventral (H) views. Teeth in wild type larvae at this stage are restricted anteriorly to the premaxillaries and dentaries of the jaw margins (A, B), and posteriorly to upper pharyngeal toothplates and the (lower) fifth ceratobranchials (C, D). Dorsal ectopic teeth in *pEF1α:EDA*-injected individuals are indicated with yellow arrows and ventral ones with arrowheads. Ectopic teeth appear in the region of the parasphenoid (arrows immediately anterior and posterior to the eye in E and F), the basibranchials (five arrowheads in E and four in midline of G) and the second through fourth ceratobranchials (arrowheads in H). One individual is represented in A-D, one in E and G, and one in F and H. Abbreviations: CB, ceratobranchial; DEN, dentary; PMX, premaxillary; PSPH, parasphenoid; UPTP, upper pharyngeal tooth plate.

### Ectopic palatal teeth in eda-overexpressing A. mexicanus

Most of the bones on which teeth have reappeared in characiform evolution are not present in 6 dpf larvae and the description of dentition is largely restricted to adult specimens. Therefore, in order to compare the dentition of *eda*-overexpressing *A*. *mexicanus* with that of other characiform species, we examined specimens at juvenile stages (25-274 dpf) in which all adult ossifications are present. We focused our analysis on the palate, both for its accessibility, as well as the fact that many of the likely reappearances of teeth in characiform evolution occurred in this region. The specific bones we scored for the presence of teeth were the palatine, ectopterygoid, endopterygoid, and metapterygoid bones of the suspensorium and the vomer and parasphenoid bones of the ventral braincase. None of these bones is toothed in wild type *A*. *mexicanus* (Valdéz-Moreno & Contreras-Balderas 2003).

We found ectopic teeth in 51 of 195 (26.2%) of juveniles injected with the *eda* expression construct and none of the 25 control *gfp*-injected juveniles (p = 0.0017, Fisher’s exact test). These ectopic teeth were located on the ectopterygoid (Fig. 4E, F, H) (n = 35 individuals, 17.9%), endopterygoid (Figure 4H) (n = 12, 6.2%), the boundary between the ectopterygoid and endopterygoid (Figure 4G) (n = 4, 2.1%), the parasphenoid (Fig. 5B) (n = 3, 1.5%) and the vomer (Fig. 5E, F) (n = 3, 1.5%). No teeth were found on the palatine (lacking teeth in all characiforms) or metapterygoid (toothed in a few characiform lineages). Interestingly, the rank order ectopterygoid > endopterygoid > parasphenoid, vomer parallels the frequency of these teeth in the order Characiformes (with parasphenoid and vomerine teeth being completely absent).

**Figure 4.**
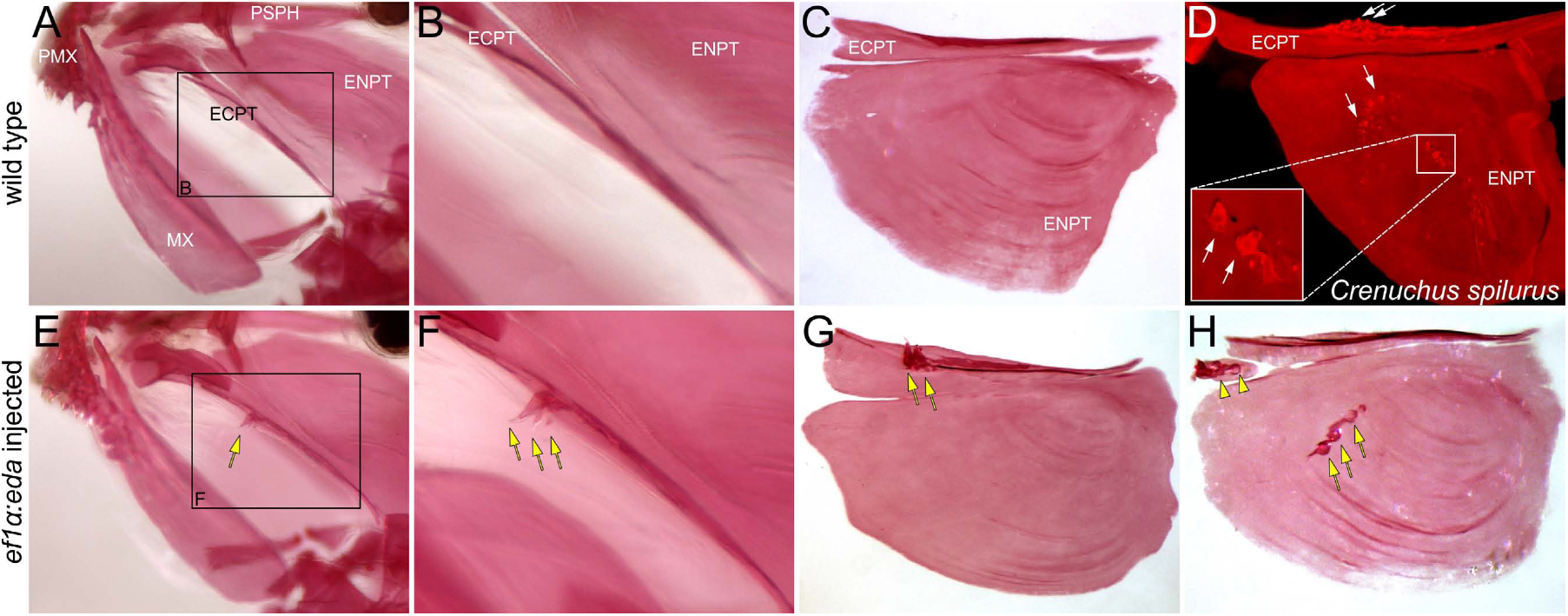
Ectopic expression of eda induces teeth on the ectopterygoid and endopterygoid of adult *A*. *mexicanus*. Cleared and alizarin-stained wild type (A-C) and *pEF1α:EDA*-injected *A*. *mexicanus* (E-H) in left lateral (A-B, E-F) and ventral (C, G-H) views. Teeth may be present on either the ectopterygoid (arrows in E-G), the endopterygoid (arrows in H), or an ectopic bone anterior to the ectopterygoid (arrowheads in H) in *pEF1α:EDA*-injected specimens but are absent from both bones in wild type *A*. *mexicanus* (A-C). Teeth (arrows in D) are present on the ectopterygoid and endopterygoid of wild type *Crenuchus spilurus* (Characiformes: Crenuchidae) for comparison. Abbreviations: ECPT, ectopterygoid; ENPT, endopterygoid; MX, maxillary; PMX, premaxillary; PSPH, parasphenoid.

**Figure 5.**
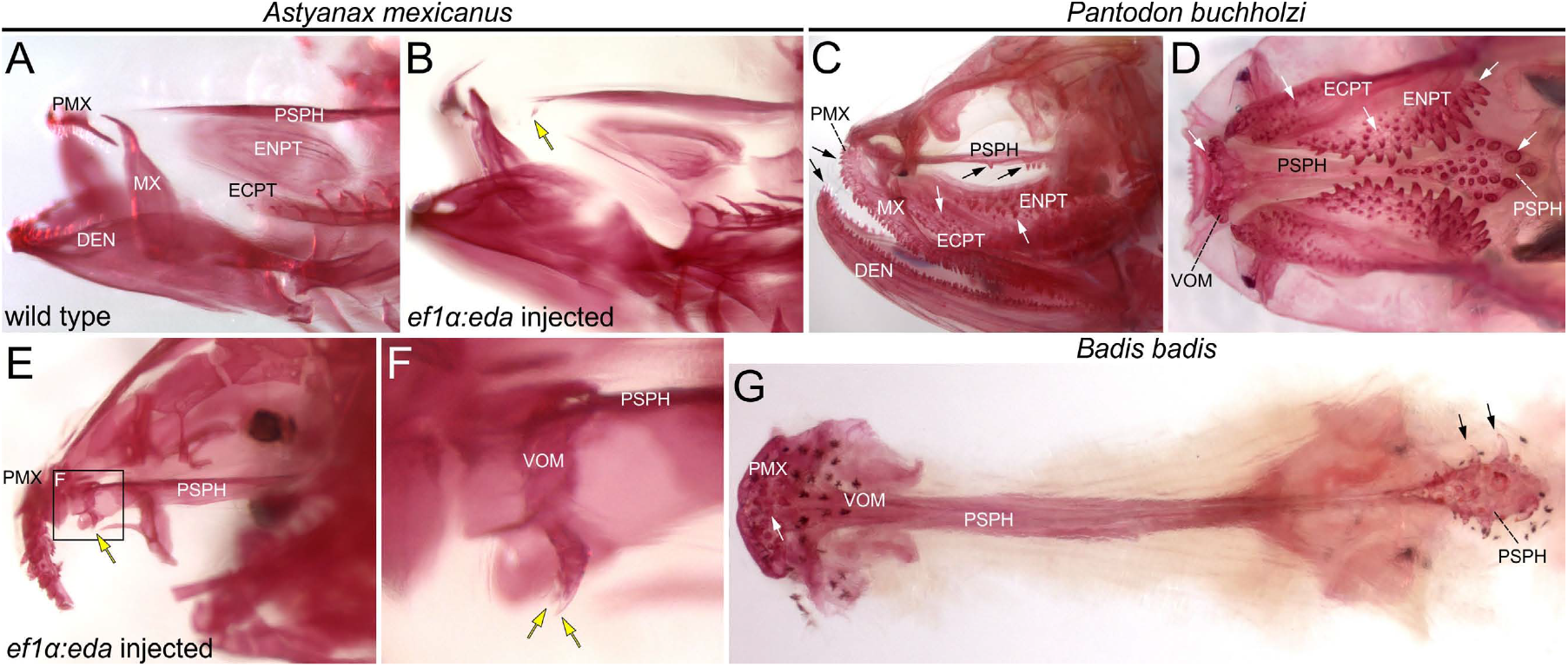
Ectopic expression of eda induces teeth on the parasphenoid and vomer of adult *A*. *mexicanus*. Cleared and alizarin-stained wild type (A) and *pEF1α:EDA*-injected *A*. *mexicanus* (B, E-F)) in left lateral. Teeth may be present on either the parasphenoid (arrow in B) or the vomer (arrows in E-F) in *pEF1α:EDA*-injected specimens but are absent from both bones in wild type *A*. *mexicanus* (A). C-D) Parasphenoid and vomerine teeth are ancestrally present in teleost fishes such as *Pantodon buccholzi* (Osteoglossiformes: Pantodontidae). Teeth in additional locations are indicated by arrows. G) According to our ancestral state reconstructions (Figure S4), parasphenoid (black arrows) and vomerine (white arrow) teeth have re-evolved in the lineage leading to *Badis badis* (Perciformes: Badidae). Abbreviations: DEN, dentary; ECPT, ectopterygoid; ENPT, endopterygoid; MX, maxillary; PMX, premaxillary; PSPH, parasphenoid; VOM, vomer.

## Discussion

### Contrasting patterns of dental diversification in cypriniforms and characiforms

Cypriniforms and characiforms are members of the Superorder Ostariophysi (Nelson *et al.* 2016), which has been considered one of nine exceptional radiations in the history of jawed vertebrates (Alfaro *et al.* 2009). Within this radiation, both groups have diversified to fill similar trophic niches, with changes in tooth shape and organization thought to have played a major role in the process (Howes 1991; Guisande *et al.* 2012, Burns & Sidlauskas 2019). The pattern of dental diversification is strikingly different between the two groups, however (Gosline 1973). Cypriniforms lack teeth in the mouth cavity but exhibit extensive variation in shape, number, and arrangement of teeth in the pharynx (Sibbing 1991; Stock 2007; Pasco-Viel et al. 2010). Variation in tooth location is limited to the simple presence (most cypriniforms) or absence (Gyrinocheilidae – algae eaters) of teeth on the fifth ceratobranchial bones of the lower posterior pharynx (Stock 2007; Nelson *et al.* 2016). Characiforms also exhibit extensive variation in tooth shape, number, and arrangement (Guisande *et al.* 2012), but this variation is largely limited to the teeth of the oral jaw margins – those of the pharynx have a simple conical shape in most groups (Roberts 1969). In addition, and in contrast to cypriniforms, characiforms exhibit substantial variation in tooth location, particularly on bones of the palate (Weitzman 1962; Roberts 1969; 1973; Fink & Fink 1981; Weitzman & Fink 1985; Toledo-Piza 2000; 2007; Novakowski *et al.* 2004; Oyakawa & Mattox 2009).

While diversification of tooth location in characiforms could simply be the result of loss from a more extensive ancestral dentition, it has been suggested that expansion of dentition has also occurred within the group, particularly for the ectopterygoid and endopterygoid teeth of the palate (Roberts 1973; Weitzman & Kanazawa 1976). We used phylogenetic character mapping to confirm these early proposals that were not based on explicit phylogenetic hypotheses. Specifically, we found evidence that ectopterygoid teeth, which were likely to have been present in the common ancestor of characiforms, were regained at least four times after being lost, while endopterygoid teeth, likely absent in this common ancestor, were gained at least five times (Fig. 2, S1, S3).

### Potential functional explanations for the regain of palatal teeth in characiform but not cypriniform evolution

An ancestral condition from which both cypriniform and characiform dentitions likely evolved is the presence of teeth throughout the oral and pharyngeal cavities (Gosline 1973). Palatal teeth in such predatory forms, represented by the modern day *Elops* (Ladyfish), serve to grasp struggling prey and facilitate its transport posteriad toward the pharynx and esophagus (Gosline 1973). A common trend in the evolution of the teleost fish dentition is its reduction in the central portion of the oral and pharyngeal cavities (including the palate) and its concentration anteriorly in oral and posteriorly in pharyngeal jaws (Gosline 1985). In characiforms, this trend is manifest in many species through specialization of oral jaw dentition for biting and shearing (Gosline 1973). A notorious example is provided by piranhas (Serrasalmidae), in which blade-like teeth allow biting pieces from animals too large to ingest. Interesting, ectopterygoid teeth, which occur in some members of this family, exhibit a similar flattened shape to the teeth of the oral jaw margins (Roberts 1969) and may also function as part of the shearing bite. Palatal teeth in other characiforms are simple cones in shape (Figure 1H, 4D) (Roberts 1969), and as they appear to be limited to insect and fish-eating species (Roberts 1973), are likely to serve the ancestral function of gripping and transporting prey.

The feeding apparatus of cypriniforms has evolved in a quite different direction from that of characiforms (Gosline 1973; Sibbing *et al.* 1986; Sibbing 1991). The posterior pharynx constitutes a powerful apparatus for mastication of food items, with teeth on hypertrophied lower pharyngeal bones biting against a dorsal keratinized pad braced by the basioccipital bone of the braincase. The mouth has become specialized for suction feeding, with a protrusible upper jaw (premaxilla) that serves a variety of functions, such as controlling the direction of water flow (Sibbing *et al.* 1986). Because teeth are absent from the mouth, palate, and upper pharynx, the function of transporting food has been assumed by muscular (dorsal) palatal and (ventral) postlingual organs that provide a peristaltic action sufficient to transport small food particles posteriorly to the masticatory apparatus (Sibbing *et al.* 1986). It is thought that the cypriniform feeding apparatus is particularly effective for feeding on plant and animal matter in bottom deposits (Gosline 1973; Sibbing *et al.* 1986; Sibbing 1991), and in such a role, palatal teeth might serve no useful role. Several lineages of cypriniforms have secondarily evolved the habit of feeding on other fishes, however, with modifications to the typical cypriniform condition including reduction of premaxillary protrusion to allow a firm grip on prey between the oral jaws (Gosline 1973; Sibbing 1991), reduction of the palatal and postlingual organs (Doosey and Bart 2011) and specialization of the pharyngeal teeth for laceration and transport of prey (Sibbing 1991). A number of authors have speculated that fish-eating cypriniforms might be more efficient predators with more extensive dentition (Nichols 1930; Weisel 1962; de Graaf et al. 2000; 2008) and indeed, radiation of such forms has occurred only in situations lacking competitors that retain oral teeth (de Graaf *et al.* 2000; 2008). We therefore suggest that the absence of palatal teeth in cypriniforms is not simply the result of absence of selection for them.

### Retention of ancestral developmental potential for palatal dentition in characiforms but not cypriniforms

Well before methods existed to test their hypothesis, Weitzman and Kanazawa (1976) proposed that “teeth and bony tooth patches remain a genetic potential for nearly any oral surface in characoids [characiforms].” We have demonstrated the existence of such potential through the overexpression of *eda* in the characiform *Astyanax mexicanus*. Some of this potential has been realized in characiform evolution in the form of reappearance of teeth on the ectopterygoid and endopterygoid bones (Fig. 6). We also found that *eda* is capable of inducing teeth in locations that are toothed in some teleosts but not in any characiform, namely the vomer and the parasphenoid on the midline of the oral cavity (Fig. 5, 6). Interestingly, vomerine teeth are likely to have reappeared in the evolution of the ostariophysan order Siluriformes (catfishes) (Fink & Fink 1981), and in an analogous situation, have been discovered as an atavism in a single individual of the Black Drum, *Pogonias cromis* (Cione & Torno 1987). It has also been suggested that teeth on the parasphenoid have reappeared in the evolution of spiny-rayed fishes (Fig. 5) (Gosline 1985); we have provided more explicit evidence that this is the case by mapping parasphenoid teeth on a phylogeny of ray-finned fishes (Fig. S4; Table S3, Supplementary References).

**Figure 6.**
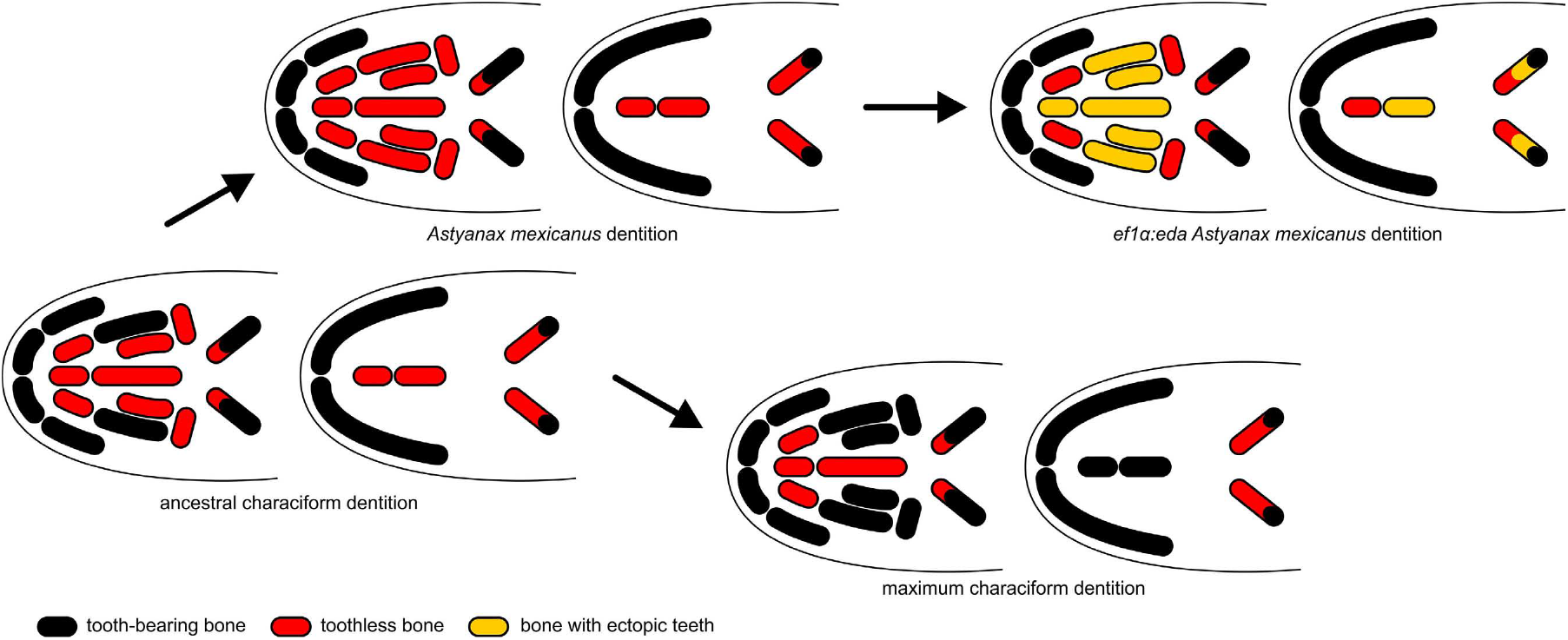
Expansion of dentition within characiforms may have been facilitated by retention of ancestral developmental potential, as seen in *A*. *mexicanus*. Schematic representation of dentition of the upper (left) and lower (right) oropharynx as in Fig. 1. The dentition of *A*. *mexicanus* is slightly reduced (loss of ectopterygoid teeth) relative to the ancestral characiform dentition (leftmost arrow). Nevertheless, this species retains the potential to form teeth in many additional locations (yellow) in response to ectopic expression of eda (upper right arrow). Some of these locations have experienced gain of teeth in characiform evolution (lower right arrow), while others have not.

The developmental potential for dentition that we have demonstrated in *Astyanax* does not appear to have been retained in the cypriniform Zebrafish. In a previous study (Aigler *et al.* 2014), we found that overexpression of ectodysplasin was capable of inducing ectopic upper pharyngeal teeth, but no teeth appeared on the palate or other regions of the oral cavity. If this restricted potential is characteristic of cypriniforms in general, it may explain the “failure” to regain palatal teeth during the radiation and trophic diversification of this group.

### The nature of the difference in retained potential for dentition between characiforms and cypriniforms

The nature of the difference in competence to produce teeth in the oral cavity between characiforms and cypriniforms remains unknown. Aigler *et al.* (2014) showed that the oral epithelium of the zebrafish retains broad competence to respond to ectodysplasin signaling with activation of NF-kappaB, a transcriptional effector of this pathway. The transcription factor *pitx2* and signaling ligand *shh* are considered markers of dental competence (Fraser *et al.* 2008) and both are present in the oral region of developing zebrafish larvae (Stock *et al.* 2006). It has been reported that *Astyanax mexicanus* has two *eda* co-orthologs, while the Zebrafish retains only one (Braasch *et al.* 2009), but how this might relate to competence to respond to the ligand with tooth initiation is unclear.

Competence to form teeth on anterior gill arches at early larval stages in *Astyanax mexicanus* might be maintained because of the later development of teeth in these locations on gill rakers (at approximately 40 dpf – Atukorala & Franz-Odendaal 2014). Not all characiforms possess such teeth and it would be interesting to determine whether species without toothed gill rakers also retain competence to form teeth in the anterior pharynx.

### Retention of ancestral developmental potential as a component of evolvability

Evolvability has been argued to be the central focus of Evolutionary Developmental Biology (Hendrikse *et al.* 2007) but what constitutes evolvability has differed widely among authors (Pigliucci 2008; Brown 2014; Payne & Wagner 2019). Features that have been proposed to contribute to evolvability include standing genetic variation in populations (Barrett & Schluter 2008; Huang 2015), key (morphological or physiological) innovations (Hunter 1998), gene or genome duplication (Ohno 1970; Cuypers & Hogeweg 2014), and features of developmental systems, such as modularity and integration (Hendrikse *et al.* 2007; Le Pabic *et al.* 2016; Fish 2019). We suggest that in the case of characiform fishes, evolvability is enhanced by retention of competence to respond to tooth induction signals in a much broader region of the oropharynx than such signals are normally produced. Such retention of ancestral developmental potential has been documented in the case of ant castes (Rajakumar *et al.* 2012); we further demonstrate that this type of evolvability differs among lineages and may have contributed to differences in morphological diversification during parallel adaptive radiations. If so, these radiations have been sculpted by “developmental push” in addition to “environmental pull” (Erwin 2017).

## Materials and Methods

### Ancestral state reconstruction

We used the character states in Mirande’s (2009) matrix of morphological features to map the presence or absence of teeth in multiple locations (premaxillaries outside of the mouth, maxillaries, ectopterygoids, endopterygoids, fourth basibranchial, gill rakers, fifth ceratobranchials, and third, fourth and fifth pharyngobranchials) on the characiform tree topology from his weighted parsimony analysis (implied weighting scheme – his Figures 1–2). Our analysis (Fig. S1, S2) was conducted with the parsimony option of Mesquite (Maddison & Maddison 2019).

Ancestral states for ectopterygoid and endopterygoid teeth were also reconstructed using the characiform molecular phylogenies of Oliveira *et al.* (2011) and Betancur-R *et al.* (2019). The tree topology and branch lengths that we used from the study of Oliveira *et al.* (2011) were based on its maximum likelihood analysis of partial sequences of two mitochondrial and three nuclear genes from 213 specimens. We assigned presence or absence of ectopterygoid teeth to 128 of these taxa and endopterygoid teeth to 94 using statements from the literature about the species, or in some cases, the genus or family to which it belonged (Table S1; Supplementary References). The Betancur-R *et al.* (2019) topology and branch lengths were from the maximum likelihood analysis of 1051 exons from 206 characiform species presented in their Figure 4. We assigned presence or absence of ectopterygoid teeth to 135 of these species and endopterygoid teeth to 126 using statements from the literature about the species or about one or more congeners (Table S2; Supplementary Reference). In both cases, the original trees were ultrametricized using the penalized likelihood with chronos command in the ape R library (Kim & Sanderson 2008) and then pruned to fit the character dataset using the drop.tip function in phytools (Revell 2012). Ancestral states were then estimated by maximum likelihood using a continuous time Markov model of binary character evolution (Mk2) with the asr.marginal function in the R package diversitree (FitzJohn 2012).

The evolution of parasphenoid teeth in ray-finned fishes was reconstructed on the relaxed molecular clock phylogeny of Farina *et al.* (2015), which they inferred from Bayesian analysis of sequences of nine nuclear genes from 285 taxa representing 284 families. We assigned presence or absence of parasphenoid teeth to each of these taxa using statements in the literature about the species or the genus, family, suborder, or order to which it belonged (Table S3; Supplementary References). When this was not possible, we used character states reported for congeneric or confamilial species. Ancestral state reconstruction was carried out as described above for characiform molecular phylogenies.

### Transient transgenic overexpression of eda in A. mexicanus

The *pEF1α:EDA* plasmid described by Aigler *et al.* (2014) contains the zebrafish *eda* coding region under the control of the *Xenopus laevis ef1α* promoter, which is expected to drive ubiquitous and continuous expression throughout development (Johnson & Krieg 1994). We modified this plasmid to allow screening injected embryos for DNA incorporation by adding an *mCherry* coding sequence with a separate *ef1α* promoter to produce *pEF1α:EDA/EF1α:mCherry*.

*Astyanax mexicanus* embryos were collected from natural spawning of laboratory populations originating from either La Cueva Chica (San Luís Potosi, Mexico) or La Cueva de El Pachón (Tamaulipas, Mexico) (Jeffery & Martasian 1998). 0.5 nl of a solution containing 30 ng *pEF1α:EDA/EF1α:mCherry* and 30 ng mRNA encoding *tol2* transposase was injected into the blastomeres of one-celled embryos. Preliminary experiments suggested that the modified plasmid produced similar results to the original *pEF1α:EDA*. Injection of *pTAL200R150G* (Urasaki *et al.* 2006), which contains an *egfp* coding region under the control of the *ef1α* promoter, served as a negative control.

### Histology

Injected individuals exhibiting mCherry fluorescence as embryos were raised to a variety of larval and juvenile stages, sacrificed, and cleared and stained for calcified structures with Alizarin red S and, in some cases, cartilage matrix with Alcian blue. These procedures followed Wise & Stock (2010) for larvae and Hanken & Wassersug (1981) for juveniles. Intact larvae were imaged in bright field with a Zeiss Axiovert 135 inverted compound microscope equipped with a Zeiss Axiocam digital camera, while juveniles were dissected before imaging in bright field or fluorescence with Zeiss Discovery V8 or Leica MZ FLIII stereomicroscopes. The former stereomicroscope was equipped with a Zeiss Axiocam MRc5 camera and the latter with a Leica DFC7000 T camera.

## Acknowledgements

We wish to thank Stacy Farina and Claudio Oliveira for sending electronic versions of phylogenetic trees from their publications and Stacy Smith for help with ancestral state reconstructions. This study was supported by grants from the US National Science Foundation (IOS-1121855 and −1755305) to DWS and from the Scientific Grant Agency of the Slovak Republic (VEGA grant No.1/0415/17) to DJ.

---

**Figure S1.**
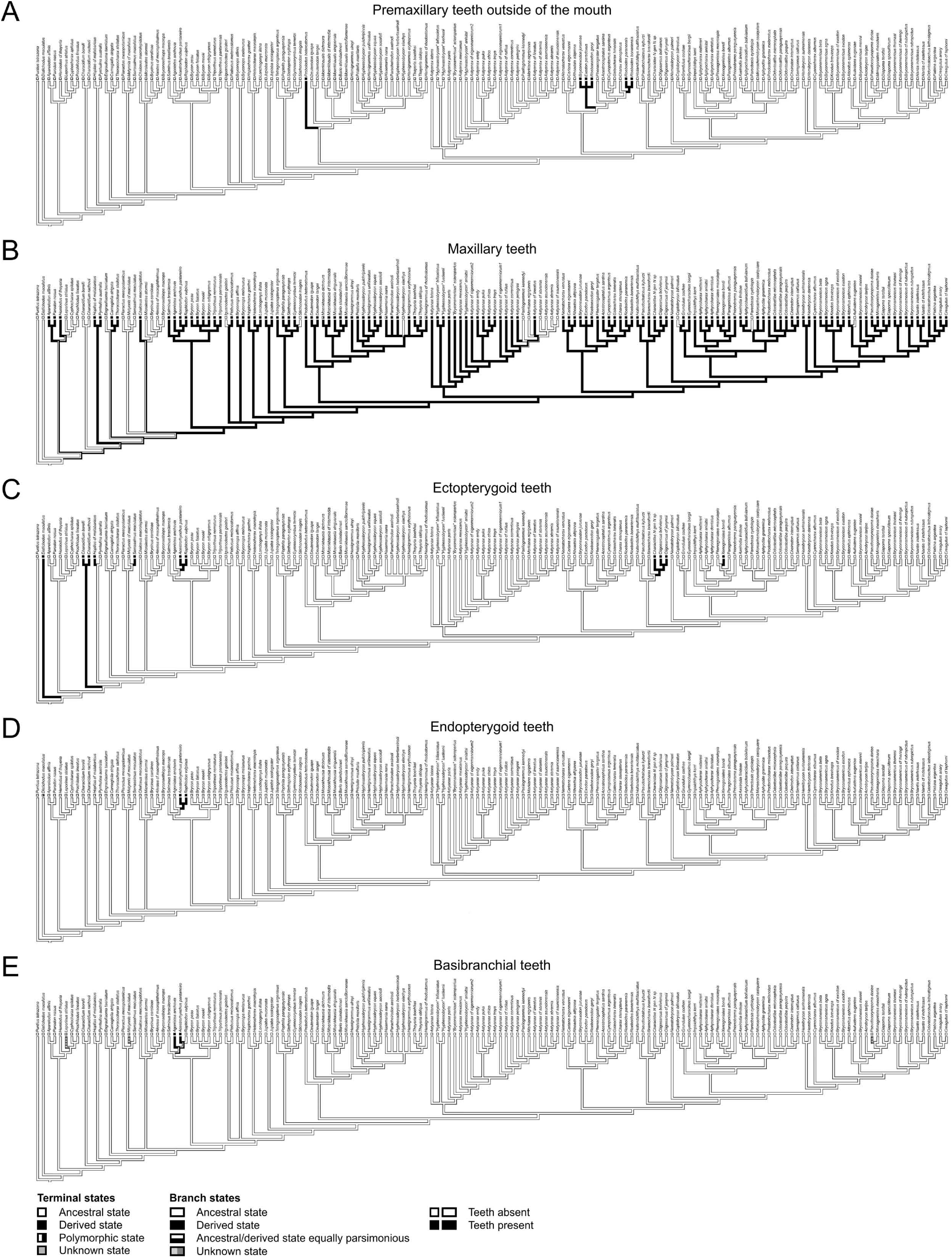
Teeth on the premaxillaries outside of the mouth, maxillaries, ectopterygoids, endopterygoids, and basibranchials have been gained in multiple characiform lineages. Parsimony reconstruction of presence (black) or absence (white) of teeth using tree topologies and character state matrices from Mirande (2009). *Puntius tetrazona* is an outgroup within the Order Cypriniformes, For the locations in this figure, presence of teeth was coded as the derived state by Mirande (2009).

**Figure S2.**
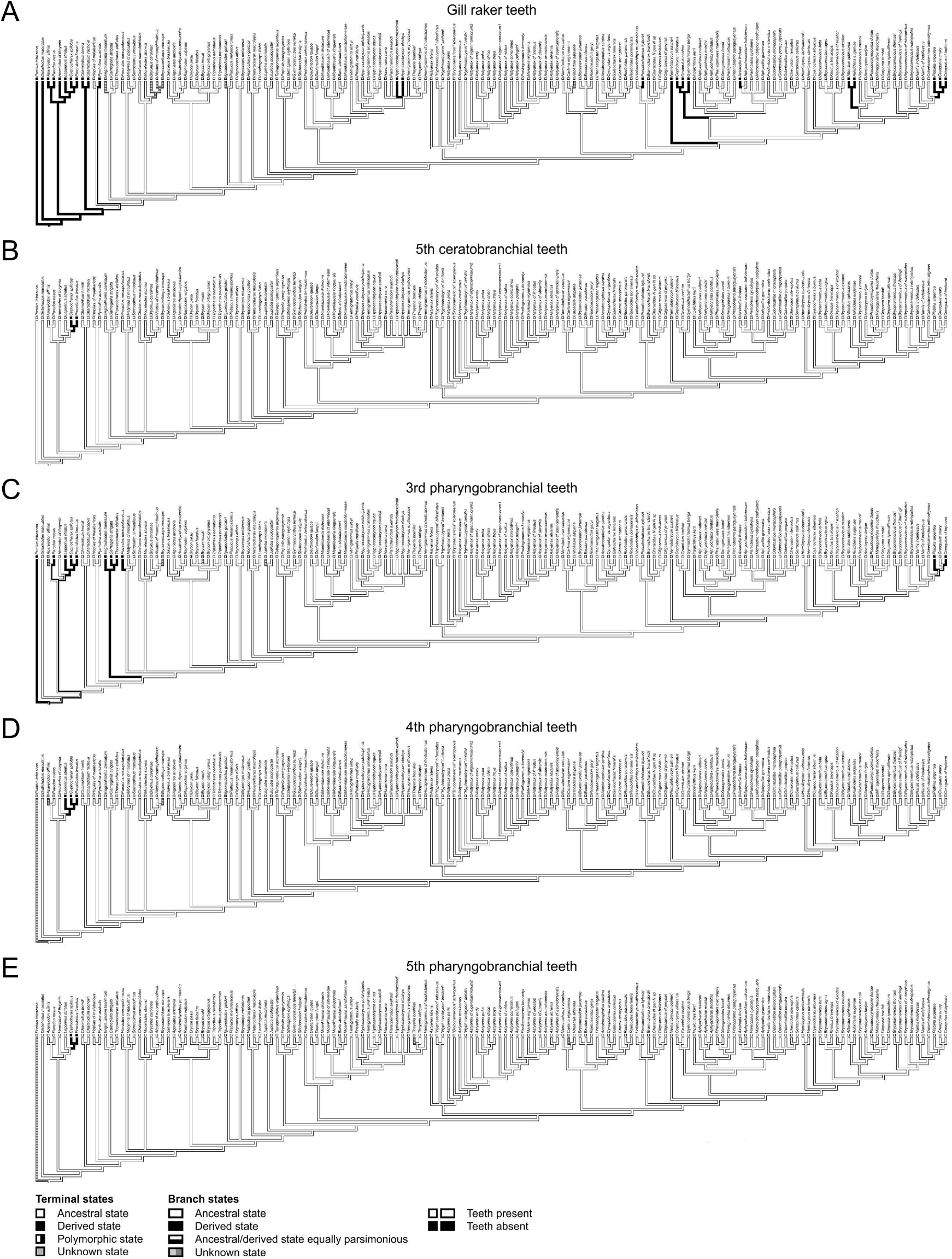
Teeth on the gill rakers have been gained within the Order Characiformes. Parsimony reconstruction of presence (white) or absence (black) of teeth using tree topologies and character state matrices from Mirande (2009). *Puntius tetrazona* is an outgroup within the Order Cypriniformes, For the locations in this figure, absence of teeth was coded as the derived state by Mirande (2009).

**Figure S3.**
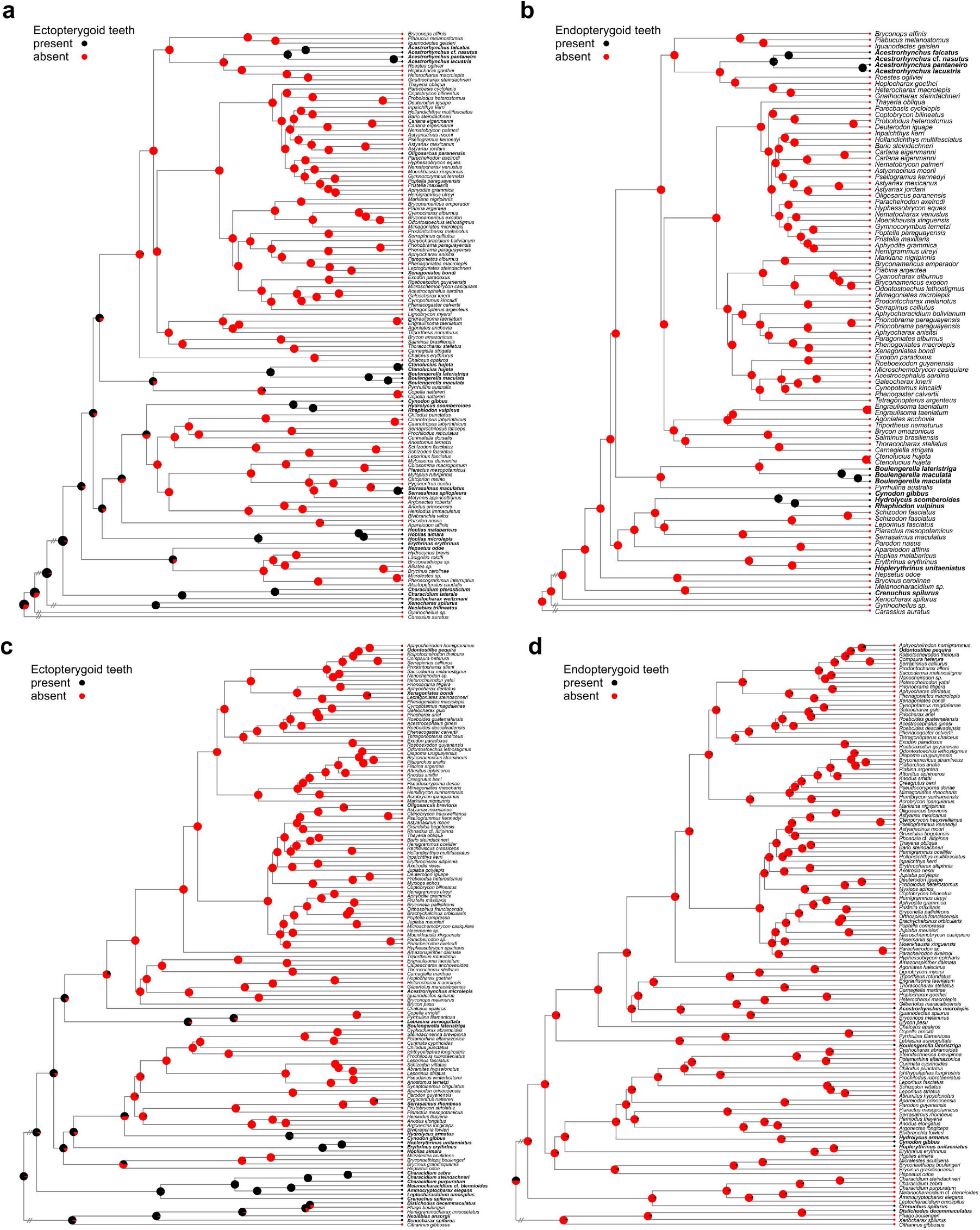
Teeth on the ectopterygoid and endopterygoid bones have been re-gained following loss in multiple characiform lineages. A-B) Trees from Fig. 2A, B depicted with full taxonomic representation. C-D) Maximum likelihood ancestral state reconstruction of presence (black) or absence (red) of teeth on the ectopterygoid (C) and endopterygoid (D) bones. Tree topologies and branch lengths are from Betancur-R *et al.* (2019) and character states were compiled from the literature (Table S1, Supplementary References). Hashmarks indicate branches not drawn to scale. Pie charts at nodes represent the relative probabilities of each character state. All taxa in S3C, D are members of the Order Characiformes. Ectopterygoid teeth (C) are reconstructed with highest probability of being present in the characiform common ancestor, with five potential regains following losses occurring within the group. Endopterygoid teeth (D) are reconstructed with equal probability of being present or absent in the characiform common ancestor, with seven potential gains occurring within the group.

**Figure S4.**
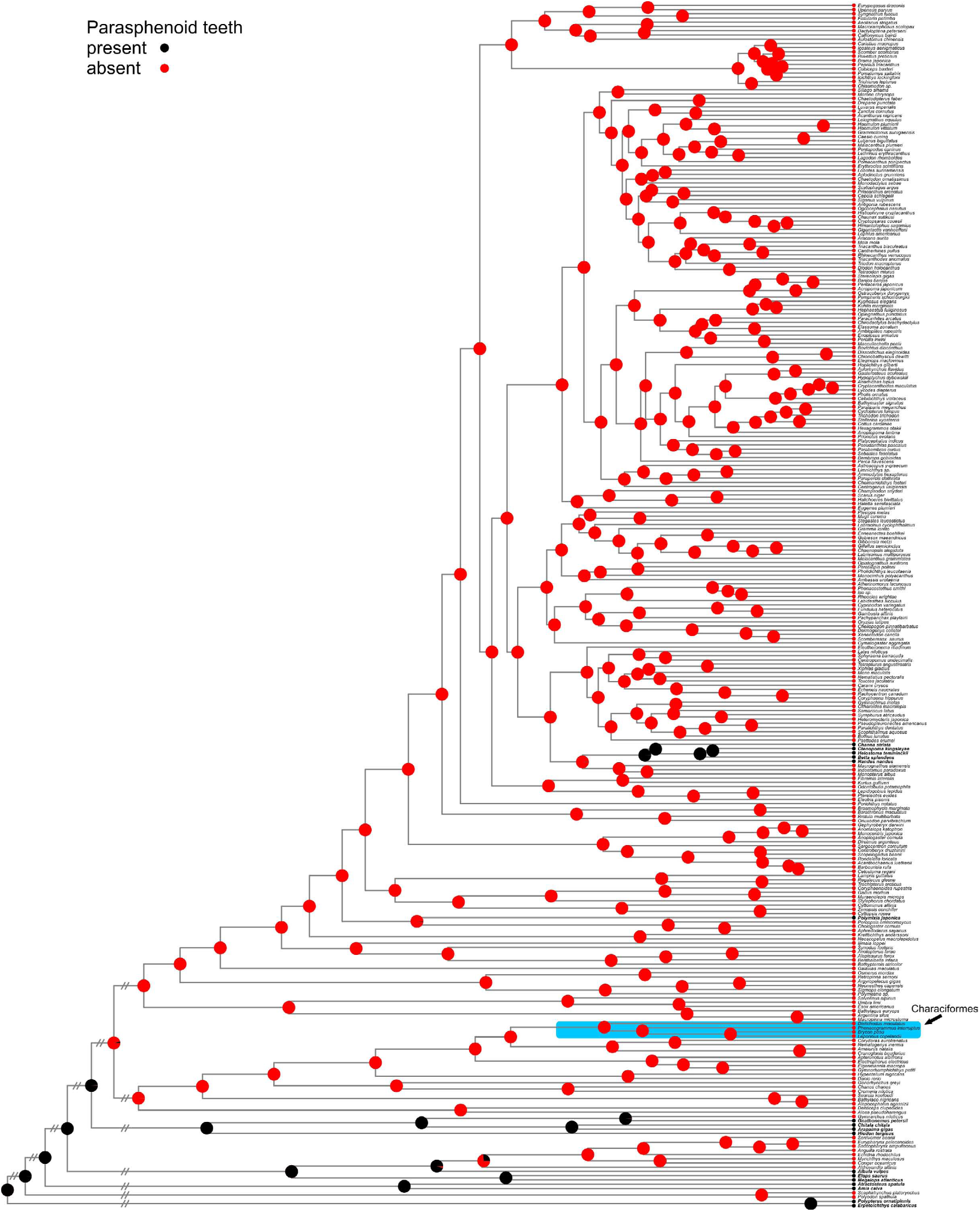
Parasphenoid teeth were present in the common ancestor of ray-finned fishes and were regained following loss twice within teleost fishes. Maximum likelihood ancestral state reconstruction of presence (black) or absence (red) of teeth on the parasphenoid bone. Tree topologies and branch lengths are from Farina *et al.* (2015) and character states were compiled from the literature (Table S3; Supplementary references). Hashmarks indicate branches not drawn to scale. Pie charts at nodes represent the relative probabilities of each character state. Lineages in which parasphenoid teeth were regained following loss are the Suborder Anabantoidei (*C*. *striata*, *C*. *kingsleyae*, *H*. *temminckii*, *B*. *splendens* and *N*. *nandus*) and the Order Polymyxiiformes (*P*. *japonica*). The order Characiformes is indicated with an arrow and blue shading.

Table S1. (not included in pdf). Ectopterygoid and endopterygoid tooth character states for Fig. 2A, B; S3A, B.

Table S2. (not included in pdf). Ectopterygoid and endopterygoid tooth character states for Fig. S3C, D.

Table S3. (not included in pdf). Parasphenoid tooth character states for Fig. S4. Supplementary References (not included in pdf)

## References

Aigler SR, Jandzik D, Hatta K, Uesugi K, Stock DW. 2014. Selection and constraint underlie irreversibility of tooth loss in cypriniform fishes. Proc. Natl. Acad. Sci. USA 111:7707–7712.

Alfaro ME, Santini F, Brock C, Alamillo H, Dornburg A, Rabosky DL, Carnevale G, Harmon LJ. 2009. Nine exceptional radiations plus high turnover explain species diversity in jawed vertebrates. Proc. Natl. Acad. Sci. USA 106:13410–13414.

Arcila D, Ortí G, Vari R, Armbruster JW, Stiassny MLJ, Ko KD, Sabaj MH, Lundberg J, Revell LJ, Betancur-R R. 2017. Genome-wide interrogation advances resolution of recalcitrant groups in the tree of life. Nat. Ecol. Evol. 1:0020.

Atukorala ADS, Franz-Odendaal T. 2014. Spatial and temporal events in tooth development of *Astyanax mexicanus*. Mech. Dev. 134:42–54.

Barrett RD, Schluter D. 2008. Adaptation from standing genetic variation. Trends Ecol. Evol. 23:38–44.

Betancur-R R, Arcila D, Vari RP, Hughes LC, Oliveira C, Sabaj MH, Ortí G. 2019. Phylogenomic incongruence, hypothesis testing, and taxonomic sampling: The monophyly of characiform fishes. Evolution 73: 329–345.

Braasch I, Brunet F, Volff JN, Schartl M. 2009. Pigmentation pathway evolution after whole-genome duplication in fish. Genome Biol. Evol. 1:479–493.

Brown RL. 2014. What evolvability really is. Brit. J. Phil. Sci. 65:549–572.

Burns MD, Sidlauskas BL. 2019. Ancient and contingent body shape diversification in a hyperdiverse continental fish radiation. Evolution 73:569–587.

Casane D, Rétaux S. 2016. Evolutionary genetics of the cavefish *Astyanax mexicanus*. Adv. Genet. 95:117–159.

Cione AL, Torno AE. 1987. Atavistic vomerine teeth in a specimen of *Pogonias cromis* (Linnaeus, 1776) (Teleostei, Perciformes). Copeia 1987:1057–1059.

Cuypers TD, Hogeweg P. 2014. A synergism between adaptive effects and evolvability drives whole genome duplication to fixation. PLoS Comput. Biol 10(04): e1003547.

de Graaf M, Dejen E, Sibbing FA, Osse JWM. 2000. The piscivorous barbs of Lake Tana (Ethiopia): major questions on their evolution and exploitation. Neth. J. Zool. 50:215–223.

de Graaf M, Dejen E, Osse JWM, Sibbing FA. 2008. Adaptive radiation of Lake Tana’s (Ethiopia) *Labeobarbus* species flock (Pisces, Cyprinidae) Mar. Freshw. Res. 59:391–407.

Doosey MH, Bart HL Jr. 2011. Morphological variation of the palatal organ and chewing pad of Catostomidae (Teleostei: Cypriniformes) J Morphol 272:1092–1108.

Erwin DH. 2017. Developmental push or environmental pull? The causes of macroevolutionary dynamics. HPLS 39:36.

Farina SC, Near TJ, Bemis WC. 2015. Evolution of the branchiostegal membrane and restricted gill openings in actinopterygian fishes. J. Morphol. 276:681–694.

Fink SV, Fink WL. 1981. Interrelationships of the ostariophysan fishes (Teleostei). Zool. J. Linn. Soc. 72:297–353.

Fish JL. 2019. Evolvability of the vertebrate craniofacial skeleton. Semin. Cell. Dev. Biol. 91:13–22.

FitzJohn RG. 2012. Diversitree: comparative phylogenetic analyses of diversification in R. Methods Ecol. Evol. 3:1084–1092.

Fraser GJ, Bloomquist RF, Streelman JT. 2008. A periodic pattern generator for dental diversity. BMC Biol. 6:32.

Gerhart J, Kirschner M. 2003. Evolvability. *In*: Keywords and Concepts in Evolutionary Developmental Biology (eds. BK Hall, WM Olson), pp. 133–137. Harvard University Press, Cambridge MA.

Gosline WA. 1973. Considerations regarding the phylogeny of cypriniform fishes, with special reference to structures associated with feeding. Copeia 1973:761–776.

Gosline WA. 1985. A possible relationship between aspects of dentition and feeding in the centrarchid and anabantoid fishes. Env. Biol. Fish. 12:161–168.

Grande L, Bemis WE. 1998. A comprehensive phylogenetic study of amiid fishes (Amiidae) based on comparative skeletal anatomy. An empirical search for interconnected patterns of natural history. J. Vert. Paleontol. 18(Suppl.):1–690.

Guisande C, Pelayo-Villamil P, Vera M, Manjarrés-Hernández A, Carvalho MR, Vari RP, Jiménez LF, Fernández C, Martínez P, Prieto-Piraquive E, Granado-Lorencio C, Duque SR. 2012. Ecological factors and diversification among Neotropical characiforms. Int. J. Ecol. 2012:610419.

Hanken J, Wassersug R. 1981. The visible skeleton. Funct. Photog. 16:22–26, 44.

Harris MP, Rohner N, Schwarz H, Perathoner S, Konstantinidis P, Nüsslein-Volhard C. 2008. Zebrafish *eda* and *edar* mutants reveal conserved and ancestral roles of ectodysplasin signaling in vertebrates. PLoS Genet. 10:e1000206.

Hendrikse JL, Parsons TE, Hallgrímsson B. 2007. Evolvability as the proper focus of evolutionary developmental biology. Evol. Dev. 9:393–401.

Hilton EJ. 2011. Bony fish skeleton. *In*: Encyclopedia of Fish Physiology: From Genome to Environment, vol. 1 (ed. AP Farrell), pp. 434–448. Academic Press, San Diego.

Howes GJ. 1991. Systematics and biogeography: an overview. *In*: Cyprinid Fishes: Systematics, Biology and Exploitation. (eds. IJ Winfield, JS Nelson), pp. 1–33. Chapman & Hall, New York.

Huang J-P. 2015. Revisiting rapid phenotypic evolution in sticklebacks: integrative thinking of standing genetic variation and phenotypic plasticity. Front. Ecol. Evol. 3:47.

Hughes LC, Ortí G, Huang Y, Sun Y, Baldwin CC, Thompson AW, Arcila D, Betancur-R, R, Li C, Becker L, Bellora N, Zhao X, Li X, Wang M, Fang C, Xie B, Zhou Z, Huang H, Chen S, Venkatesh B, Shi Q. 2018. Comprehensive phylogeny of ray-finned fishes (Actinopterygii) based on transcriptomic and genomic data. Proc. Natl. Acad. Sci USA 115:6249–6254.

Hunter JP. 1998. Key innovations and the ecology of macroevolution. Trends Ecol. Evol. 13:31–36.

Irisarri I, Baurain D, Brinkmann H, Delsuc F, Sire J-Y, Kupfer A, Petersen J, Jarek M, Meyer A, Vences, Philippe H. 2017. Phylotranscriptomic consolidation of the jawed vertebrate timetree. Nature Ecol. Evol 1:1370–1378.

Jeffery WR. 2009. Regressive evolution in *Astyanax* cavefish. Annu. Rev. Genet. 43:25–47.

Jeffery WR, Martasian DP. 1998. Evolution of eye regression in the cavefish *Astyanax*: apoptosis and the *Pax-6* gene. Amer. Zool. 38:685–696.

Johnson AD, Krieg PA. 1994. pXeX, a vector for efficient expression of cloned sequences in Xenopus embryos. Gene 147:223–226.

Kim J, Sanderson MJ. 2008. Penalized likelihood phylogenetic inference: bridging the parsimony-likelihood gap. Syst. Biol. 57:665–674.

Le Pabic P. Cooper WJ, Schilling TF. 2016. Developmental basis of phenotypic integration in two Lake Malawi cichlids. EvoDevo 7:3.

Maddison WP, Maddison DR. 2019. Mesquite: a modular system for evolutionary analysis. Version 3.61 http://www.mesquiteproject.org.

Mattox GMT, Toledo-Piza. 2012. Phylogenetic study of the Characinae (Teleostei: Characiformes: Characidae). Zool. J. Linn. Soc. 165:809–915.

Mirande JM. 2009. Weighted parsimony phylogeny of the family Characidae (Teleostei: Characiformes). Cladistics 25:574–613.

Mirande JM 2010. Phylogeny of the family Characidae (Teleostei: Characiformes): from characters to taxonomy. Neotrop. Ichthyol. 8:385–568.

Mirande JM. 2019. Morphology, molecules and the phylogeny of Characidae (Teleostei, Characiformes). Cladistics 35:282–300.

Nelson JS, Grande TC, Wilson MVH. 2016. Fishes of the World, 5^th^ ed. Wiley, Hoboken NJ.

Nichols JT. 1930. Speculation on the history of the Ostariophysi. Copeia 1930:148–151.

Novakowski GC, Fuji R, Hahn NS. 2004. Diet and dental development of three species of *Roeboides* (Characiformes: Characidae). Neotrop. Ichthyol. 2:157–162.

Ohno S. 1970. Evolution by Gene Duplication. Springer-Verlag, New York.

Oliveira C, Avelino GS, Abe KT, Mariguela TC, Benine RC, Ortí G, Vari RP, Corrêa e Castro RM. 2011. Phylogenetic relationships within the speciose family Characidae (Teleostei: Ostariophysi: Characiformes) based on multilocus analysis and extensive ingroup sampling. BMC Evol. Biol. 11:275.

Oyakawa OT, Mattox GMT. 2009. Revision of the Neotropical trahiras of the *Hoplias lacerdae* species-group (Ostariophysi: Characiformes: Erythrinidae) with descriptions of two new species. Neotropical Ichthyol. 7:117–140.

Pasco-Viel E, Charles C, Chevret P, Semon M, Tafforeau P, Viriot L, Laudet V. 2010. Evolutionary trends of the pharyngeal dentition in Cypriniformes (Actinopterygii: Ostariophysi). PLoS One 5:e11293.

Payne JL, Wagner A. 2019. The causes of evolvability and their evolution. Nat. Rev. Genet. 20:24–38.

Pigliucci M. 2008. Is evolvability evolvable? Nat. Rev. Genet. 9:75–82.

Rajakumar R, San Mauro D, Dijkstra MB, Huang MH, Wheeler DE, Hiou-Tim F, Khila A, Cournoyea M, Abouheif E. 2012. Ancestral developmental potential facilitates parallel evolution in ants. Science 335:79–82.

Revell LJ. 2012. phytools: an R package for phylogenetic comparative biology (and other things). Mol. Biol. and Evol. 3: 217–223.

Roberts T. 1969. Osteology and relationships of characoid fishes, particularly the genera *Hepsetus*, *Salminus*, *Hoplias*, *Ctenolucius*, and *Acestrorhynchus*. Proc. Cal. Acad. Sci. 36:391–500.

Roberts T. 1973. Interrelationships of ostariophysans. *In*: Interrelationships of Fishes (eds. PH Greenwood, RS Miles, C Patterson), pp. 142–157. Academic Press, New York.

Sibbing FA, Osse JWM, Terlouw A. 1986. Food handling in the carp (*Cyprinus carpio*): Its movement patterns, mechanisms and limitations. J. Zool. 210:161–203.

Sibbing FA. 1991. Food capture and oral processing. *In*: Cyprinid Fishes: Systematics, Biology and Exploitation. (eds. IJ Winfield, JS Nelson), pp. 377–412. Chapman & Hall, New York.

Stock DW. 2001. The genetic basis of modularity in the development and evolution of the vertebrate dentition. Phil. Trans. R. Soc. Lond. B 356:1633–1653.

Stock DW. 2007. Zebrafish dentition in comparative context. J. Exp. Zool. B Mol. Dev. Evol. 308:523–549.

Stock DW, Jackman WR, Trapani J. 2006. Developmental genetic mechanisms of evolutionary tooth loss in cypriniform fishes. Development 133:3127–3137.

Toledo-Piza M. 2000. The Neotropical fish subfamily Cynodontinae (Teleostei: Ostariophysi: Characiformes): a phylogenetic study and a revision of Cynodon and Raphiodon. Amer. Mus. Novitates 3286:1–88.

Toledo-Piza M. 2007. Phylogenetic relationships among *Acestrorhynchus* species (Ostariophysi: Characiformes: Acestrorhynchidae) Zool. J. Linn. Soc. 151:691–757.

Trapani J, Yamamoto Y, Stock DW. 2005. Ontogenetic transition from unicuspid to multicuspid oral dentition in a teleost fish: *Astyanax mexicanus*, the Mexican tetra (Ostariophysi: Characidae). Zool. J. Linn. Soc. 145:523–538.

Urasaki A, Morvan G, Kawakami K. 2006. Functional dissection of the *Tol2* transposable element identified the minimal *cis*-sequence and a highly repetitive sequence in the subterminal region essential for transposition. Genetics 174:639–649.

Valdéz-Moreno ME, Contreras-Balderas S. 2003. Skull osteology of the characid fish *Astyanax mexicanus* (Teleostei: Characidae). Proc. Biol. Soc. Wash. 116:341–355.

Wagner GP, Altenberg L. 1996. Perspective: Complex adaptations and the evolution of evolvability. Evolution 50:967–976.

Watson RA, Szathmáry E. 2016. How can evolution learn? Trends Ecol. Evol. 31:147–157.

Watson RA, Wagner GP, Pavlicav M, Weinreich DM, Mills R. 2014. The evolution of phenotypic correlations and “developmental memory”. Evolution 68:1124–1138.

Weisel GF. 1962. Comparative study of the digestive tract of a sucker, Catostomus catostomus, and a predaceous minnow, Ptychocheilus oregonense. Am. Midl. Nat. 68:334–346.

Weitzman SH. 1962. The osteology of *Brycon meeki*, a generalized characid fish, with an osteological definition of the family. Stanford Ichthyol. Bull. 8:1–77.

Weitzman, SH, Fink SV. 1985. Xenurobryconin phylogeny and putative pheromone pumps in Glandulocaudine fishes (Teleostei: Characidae). Smithson. Contr. Zool. 421:1–118.

Weitzman SH, Kanazawa RH. 1976. *Ammocryptocharax elegans*, a new genus and species of riffle-inhabiting characoid fish (Teleostei: Characidae) from South America. Proc Biol. Soc. Wash. 89:325–346.

Wise SB, Stock DW. 2010. *bmp2b* and *bmp4* are dispensable for zebrafish tooth development. Dev. Dyn. 239:2534–2546

